# Aging drives a program of DNA methylation decay in plant organs

**DOI:** 10.1101/2024.11.04.621941

**Authors:** Dawei Dai, Ken Chen, Jingwen Tao, Ben P. Williams

## Abstract

How organisms age is a question with broad implications for human health. In mammals, DNA methylation is a biomarker for biological age, which may predict age more accurately than date of birth. However, limitations in mammalian models make it difficult to identify mechanisms underpinning age-related DNA methylation changes. Here, we show that the short-lived model plant *Arabidopsis thaliana* exhibits a loss of epigenetic integrity during aging, causing heterochromatin DNA methylation decay and the expression of transposable elements. We show that the rate of epigenetic aging can be manipulated by extending or curtailing lifespan, and that shoot apical meristems are protected from this aging process. We demonstrate that a program of transcriptional repression suppresses DNA methylation maintenance pathways during aging, and that mutants of this mechanism display a complete absence of epigenetic decay. This presents a new paradigm in which a gene regulatory program sets the rate of epigenomic information loss during aging.

## Main Text

Aging is a complex multifactorial biological process that is observed across the tree of life (*1*). Understanding how and why living organisms age is a central question in biology and the treatment of disease. The past century has witnessed unprecedented strides in identifying drivers of the aging process and its hallmarks in mammals (*2, 3*). However, less attention has been paid to the field of plant aging, a complex process that differs fundamentally from that in animals (*4-7*). Despite conceptual debates, plant aging generally refers to functional decline that occurs with time in an individual plant and its component cells and organs (*8*).

Cytosine DNA methylation is a dynamic and heritable epigenetic mark that is essential for diverse biological processes in plants and animals (*9*). DNA methylation changes in a subset of CGs have emerged as a leading aging biomarker (known as “epigenetic clock”) for accurately estimating biological age in humans and several other mammal species (*10-12*). Mammalian epigenetic aging clocks capture age-related gains in CG methylation within CpG islands, as well as DNA methylation losses within heterochromatin (*13-16*). Currently, the relationship between aging and DNA methylation dynamics in plants, which lack CpG islands, is largely unknown. Unlike mammals, where DNA methylation predominantly occurs in the symmetric CG context, plants also catalyze cytosine methylation in all sequence contexts (CG, CHG, and CHH contexts, where H = A, T, or C) (*17*). Despite these differences, many of the major DNA methylation maintenance mechanisms are highly conserved between plants and animals, and likely evolved at the common ancestor to all eukaryotes (*18*).

While epigenetic clocks have become a widely accepted robust method of age estimation in mammals, the molecular mechanisms underpinning them remain very poorly understood (*13*). One cause of this gap in knowledge is that cell culture systems are inadequate to study organismal aging, and the complexity of mammalian model species makes rapid, high throughput study of genetic mechanisms challenging. Additionally, invertebrate animal aging models such as *Caenorhabditis elegans* lack DNA methylation (*19*). Here we show that heterochromatin regions in the genome of the short-lived model plant species *Arabidopsis thaliana* undergoes a pattern of epigenetic decay that is similar to DNA methylation losses that accumulate during human and mammalian aging, as well as in cancer (*20-24*). We demonstrate that rates of epigenetic decay can be modulated by altering growth conditions and genetic manipulation of plant aging pathways, strongly suggesting that methylation patterns reflect an underlying “biological age” that can be decoupled from calendar age. We also show that the shoot apical meristem is protected from epigenetic aging, and that new organs emerge as epigenetically “young”, even in older plants. Finally, we show that a mechanism of transcriptional repression underpins the age-related loss of DNA methylation in heterochromatin, and that abolishing this mechanism creates plants with permanently “young” epigenomes. This finding represents a new mechanistic insight into epigenetic aging and suggests that DNA methylation decay may be a regulated program that could potentially be modulated to alter epigenetic aging clocks in eukaryotes.

### Heterochromatin DNA methylation decay during aging in *A. thaliana*

We set out to examine if plants undergo changes to the methylome during aging as has been established in mammals. Under long-day growth conditions (LD, 16 h light/ 8 h dark), the first true leaves of *A. thaliana* (accession Col-0, hereafter termed wild-type [WT]) undergo a complete life cycle consisting of a series of developmental and senescent processes within about 7 weeks (*25*) (Fig. 1A). To quantitatively define the aging stages of these leaves, we performed qRT-PCR analysis using marker genes corresponding to each key developmental stage: *CYCB1;2* for cell proliferation (*26*), *PIF4* for cell expansion (*27*), *FT* for the reproductive transition (*28, 29*), and *ORE1* for senescence (*30*) (Fig. 1B). We next generated single baseresolution methylomes for the first true leaves from various calendar ages (Fig. 1A) using Enzymatic Methyl sequencing (EM-seq) (*31*) (Table S1). Compared to embryos (*32*) (Table S2), aging leaves displayed a progressive age-dependent reduction of CG methylation (mCG) within pericentromeric regions (Fig. 1C and fig. S1. A and B). We did not observe similar dynamics for CHG and CHH methylation (Fig. S1B). In plants, pericentromeric regions form heterochromatin dominated by transposable elements (TEs) and other repetitive sequences, whereas chromosome arms are gene-enriched and predominantly euchromatic (Fig. S1C). We found that mCG level progressively decreased with age over all TEs, but to a much lesser extent over methylated genes (*33, 34*) or their adjacent intergenic sequences (Fig. 1D and fig. S2). These observations suggest that age-dependent loss of mCG predominantly occurs in repetitive heterochromatin regions. In plant somatic tissues, mC can be removed enzymatically by a group of demethylases (collectively named DRDD), which predominantly function in gene-rich regions (*35-38*). The genes encoding DRDD demethylases are expressed during leaf development and maturation (Fig. S4A) and could therefore feasibly underpin the loss of heterochromatin mCG during aging. However, we observed no differences in heterochromatin methylation patterns between WT (WT segregants for *drdd*) and triple (*rdd*) or quadruple (*drdd*) mutant methylomes of the same age (*35*) (Fig. S4, B and C), suggesting that active demethylation by DRDD does not contribute to age-related methylation losses.

**Fig. 1.**
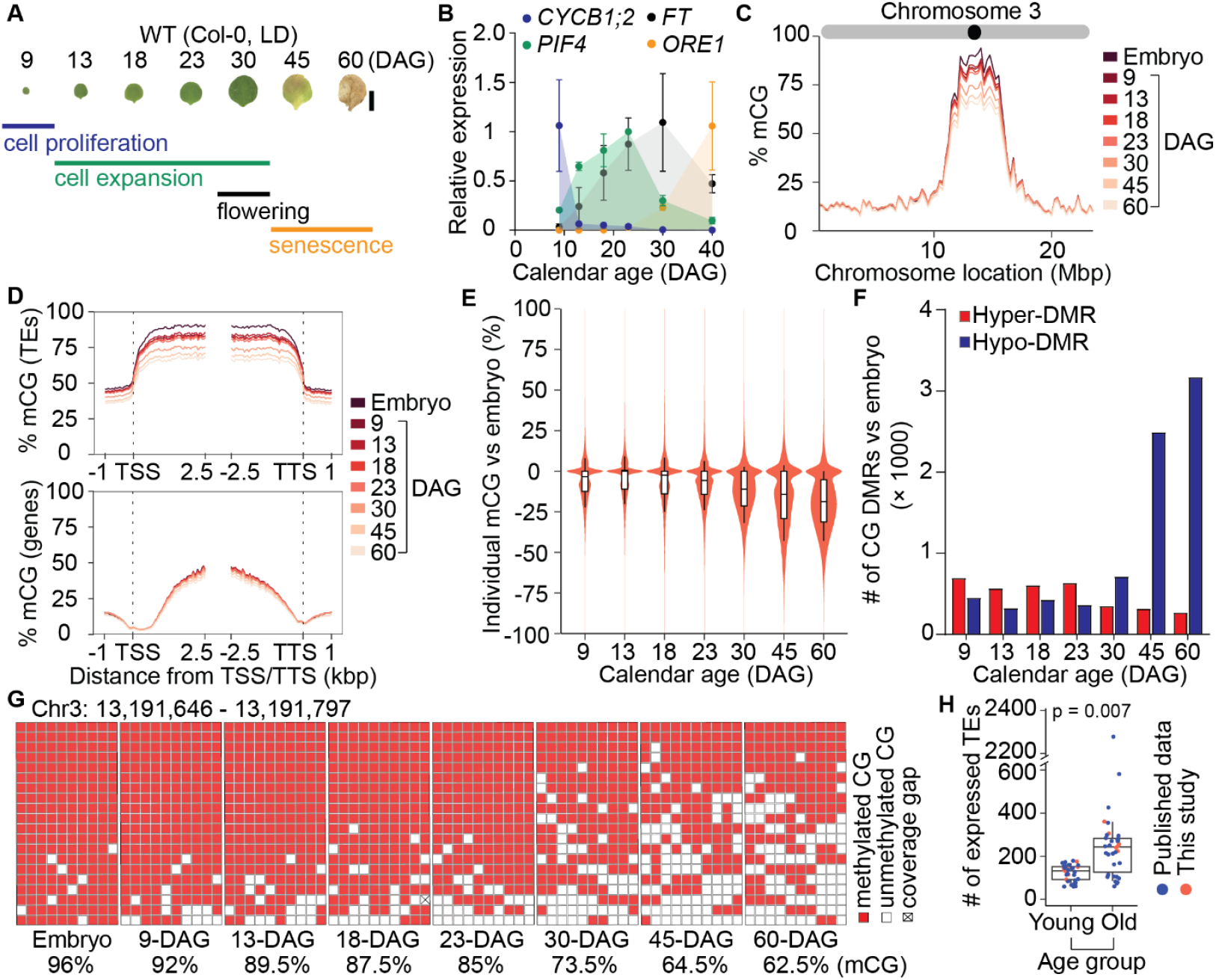
DNA methylation dynamics during leaf aging in *A. thaliana*. **(A)** The aging process of WT first true leaves in long day (LD, 16 h light/ 8 h dark) conditions. Key developmental stages of cell proliferation, cell expansion, flowering, and senescence were defined based on the expression patterns of marker genes in **(B)**. Scale bar = 0.5 cm. **(B)** Quantitative RT-PCR analysis of *CYCB1;2, PIF4, FT*, and *ORE1* transcript levels in aging first true leaves, across three biological replicates for each time point. Error bars represent standard deviations of the mean from three biological replicates. The values of each gene were relative to its highest expression (set as 1). **(C)** Average chromosome 3 mCG levels across 250 kbp windows in embryo and aging first true leaves. **(D)** Distribution of mean % mCG across total transposable elements (TEs) and genes and the flanking 1-kbp regions in embryo and aging first true leaves. **(E)** Violin plot showing the difference in percent methylation of individual CGs (within TEs) between embryo and aging first true leaves. Boxes bound the interquartile range, lines represent the median, and whiskers denote the 10th and 90th percentiles. **(F)** Number of CG DMRs in aging first true leaves compared to embryo. **(G)** Brick plot of a representative TE fragment with ten CGs covered by continuous read pairs. Each column represents an individual CG and each row represents an independent read pair. **(H)** The number of expressed TEs in young (<= 28-DAG) vs. old (>28-DAG) rosette leaves. Statistical significance was determined using one-tailed Welch’s t-test.

To assess the impact of aging on individual mCG dynamics, we quantified mCG differences in a pairwise fashion between aging first true leaves and embryos. We observed a progressive increase in the number of hypo-cytosines (decreased mCG) over TEs with age (Fig. 1, E and G and fig. S3B), but not genes (Fig. S3, A and B). We then calculated differentially methylated regions (DMRs) in aging first true leaves compared with the embryo. A sharp increase in the number of hypo-DMRs located within pericentromeric regions (Fig. S3C) was observed in first true leaves older than 30-DAG (Fig. 1F and table S3), indicating that cumulative single-mCG losses during aging add up to major region-scale losses of mCG during senescence. To examine whether such methylation losses meaningfully impact transcriptional silencing and genome defense in repetitive DNA, we analyzed the number of expressed full-length TEs across all ages in our data (Table S4) and published studies (Table S5) of transcriptome dynamics during leaf aging. In both cases, there was a clear age-dependent upregulation of TEs, with a greater number of TEs identified as transcriptionally active in older vs. younger tissues (Fig. 1H, fig. S3D, and table S6).

To assess whether patterns of individual mCG decay during aging could serve as an accurate predictor of biological age, analogous to epigenetic aging clocks that have been established in mammals (*10, 39*), we obtained and analyzed 94 published methylomes of WT leaf and seedling tissues from 48 studies that used conventional (though not identical) growth conditions and sampling methods (Table S2). For all studies, the calendar age of samples was recorded in the methodology, except for five studies in which age was recorded with specific developmental information (e. g., bolting, late senescence). For these samples, we estimated calendar ages based on the timing of these developmental stages in our growth conditions (Table S2). We then performed a linear regression of average mCG within TEs over calendar age for all 101 methylomes (Table S7). This regression revealed a significant and robust relationship between calendar age and methylation level (p = 1e-17) that explains over half of the variance in the dataset (R^2^ = 0.526) (Fig. 2A).

**Fig. 2.**
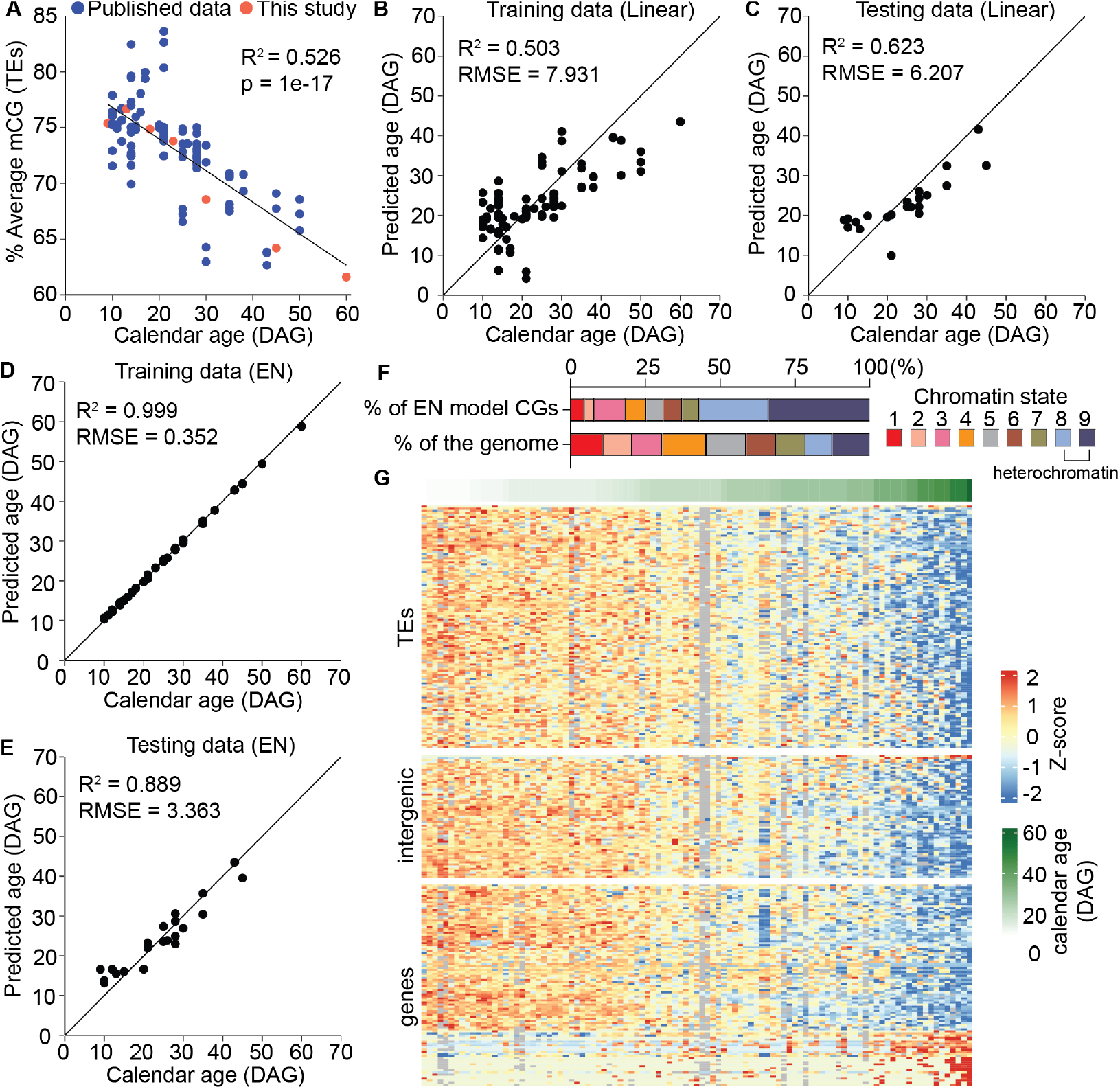
DNA methylation is a robust predictor of calendar age in *A. thaliana*. **(A)** Linear regression showing the relationship between reported calendar age and % mCG across all TEs in 101 methylomes. **(B)** Predictive accuracy of linear regression between calendar age and % mCG within TEs on a randomly selected training dataset of 80 methylomes. **(C)** Predictive accuracy of linear regression between calendar age and % mCG within TEs on 21 methylomes selected as a testing dataset. **(D)** Predictive accuracy of a trained elastic net regression model training data. **(E)** Predictive accuracy of a trained elastic net regression model on testing data. **(F)** Distribution of 276 EN-identified CG sites within chromatin states. **(G)** Heatmap showing mCG levels of 276 EN-identified CG sites across 101 methylomes. Columns are ordered by age. Heatmap values are Z-scores.

To achieve greater predictive accuracy, we trained a machine learning elastic net (EN) regression model to construct an epigenetic aging clock (*11, 40-42*). The 101 methylomes were split into randomly selected training (80 samples) and testing (21 samples) sets (Table S8) to train an EN model (see Supplemental Methods), which identified 276 CG sites as high confidence predictors for calendar age (Table S9). When compared with the linear regression model using average methylation within TEs, the EN regression model had a markedly smaller RMSE on the testing data set (3.363 days vs 6.207 days) (Fig. 2, B to E). We observed that the majority of CGs identified by the EN model are situated in TEs and intergenic regions, as well as preferentially localized within heterochromatin (defined by (*43*)) (Fig. 2F). The EN model therefore mostly identified similar DNA methylation dynamics to our analysis of methylation levels within TEs (Fig. 2, D and E), with the majority of CGs exhibiting methylation declines during leaf aging (Fig. 2G). Consistently, we found that an EN model trained only on TE methylation patterns predicted age almost as accurately as the model trained on whole genome data (3.805 days vs 3.363 days RMSE) (Fig. S5). A small subset of CGs within genes gain methylation during aging, not unlike the gains of methylation within CpG islands that are a hallmark of epigenetic aging clocks in mammals (*14*). Due to its ability to capture more complex relationships between specific sites and aging patterns, the EN model therefore represents a capable predictor of calendar age based on genomic mCG patterns.

### Different aging rates lead to divergent methylomes

An accurate biological aging predictor should be able to reflect instances where the underlying biological age is decoupled from the calendar age of the individual. For example, in humans, if individuals age at different rates, their methylomes are likely to become more distinct over time (*44*). Likewise, epigenetic clock models detect different aging rates in age-related diseases such as progeria (*45*), and after treatments that alter lifespan in mice (*46*). Due to their flexible and environmentally responsive developmental programs, plants offer unique opportunities to decouple the rate of biological aging from calendar age. We leveraged short-day (SD, 8 h light/ 16 h dark) growth conditions that extensively prolong lifespan and organ longevity to test the impact on rates of epigenetic decay (Fig. 3, A and B, and fig. S6A), generating methylomes for SD-grown first true leaves of various age stages (Table S1). While TE mCG levels were similar between SD and LD plants in the youngest leaves, they diverged after 23-DAG (Fig. 3, C and D, fig. S6, B to D, and table S7), showing a substantially slower rate of epigenetic decay in the slower-aging SD plants (Fig. 3E and table S10), consistent with their longer lifespan. We also obtained and analyzed public methylomes of rosette leaves from SD-grown plants (Table S2) and found a similar divergence compared to LD plants (Fig. S6D and table S7). Consequently, our data captures rates of epigenetic decay that reflect rates of biological aging and lifespan.

**Fig. 3.**
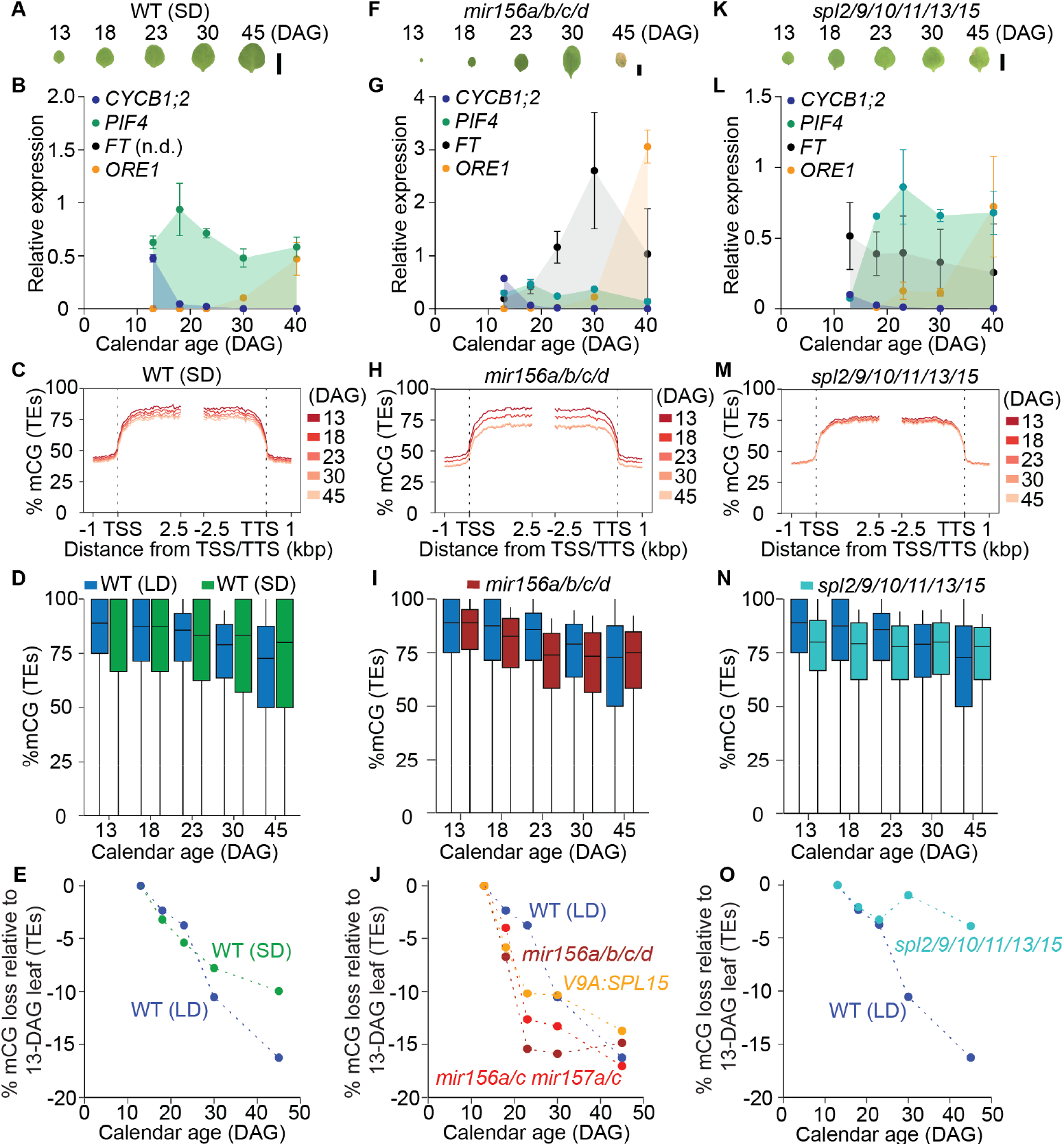
Different biological aging rates lead to divergent methylomes. **(A, F**, and **K)** First true leaf phenotypes of WT in short day (SD, 8 h light/ 16 h dark) conditions **(A)**, *mir156a/b/c/d* **(F)**, and *spl2/9/10/11/13/15* **(K)** in LD conditions. Bars = 0.5 cm. **(B, G**, and **L)** Quantitative RT-PCR analysis of *CYCB1;2, PIF4, FT*, and *ORE1* transcript levels in aging first true leaves of WT (SD) **(B)**, *mir156a/b/c/d* **(G)**, and *spl2/9/10/11/13/15* **(L)**. Error bars represent standard deviations of the mean from three biological replicates. The values of each gene were relative to its highest expression in WT (LD) aging first true leaves (see Fig. 1). **(C, H**, and **M)** Distribution of mean % mCG over TEs and the flanking 1-kbp of WT (SD) **(C)**, *mir156a/b/c/d* **(H)**, and *spl2/9/10/11/13/15* **(M). (D, I**, and **N)** Boxplot showing comparisons of mCG level in aging first true leaves between WT (LD) and WT (SD) **(D)**, *mir156a/b/c/d* **(I)**, and *spl2/9/10/11/13/15* **(N)**. Boxes represent the interquartile range, horizontal lines denote the median, and whiskers denote the 10th and 90th percentiles. All pairwise differences between WT (LD) and WT (SD) or mutants are significant (Mann-Whitney U test, p = <0.0001), with the exception of 45-DAG *mir156a/b/c/d*. **(E, J**, and **O)** Comparison of the rate of mCG decay in first true leaves between WT (LD) and WT (SD) **(E)**, *mir156a/b/c/d* **(J)**, and *spl2/9/10/11/13/15* **(O)**.

To further demonstrate that epigenetic decay reflects underlying biological aging, we analyzed methylomes of genetic mutants with altered aging processes. The *microRNA156/157* (*miR156/157*) family and its target SQUAMOSA PROMOTER BINDING PROTEIN-LIKEs (SPLs) transcription factors define a key developmental process by regulating the juvenile:adult transition, which strongly influences organ maturation and lifespan in plants (*47, 48*). We generated a time-course of aging first true leaf methylomes from two obtained quadruple mutants, *mir156a/b/c/d* and *mir156a/c mir157a/c* (*49*), and a *miR156*-resistant *SPL15* overexpression line *V9A:rSPL15*, which display accelerated development with a truncated juvenile stage (*50*) (Fig. 3, F and G, and fig. S7). Unlike WT, where mCG decreased progressively with age, these mutants exhibited dramatic loss of mCG at 18- and 23-DAG over TEs (Fig. 3, H and I, fig. S7, and table S7), coinciding with the typical timing of the juvenile:adult phase transition. Consequently, the rate of mCG decay was significantly increased in all mutants compared with that in WT (LD) (Fig. 3J and table S10). Accelerated epigenetic aging has also been observed at reproductive transitions in mammals (*51*), suggesting that reproductive investment accelerates epigenetic aging in diverse lineages.

We next tested a longer-lifespan hextuple mutant *spl2/9/10/11/13/15* that shows delayed vegetative phase change (*52*) (Fig. 3, K and L, and fig. S8A). SPLs (which are targeted for degradation by *miR156/157*) regulate aging in the opposite way to *miR156/157*, by promoting the juvenile:adult transition in *A. thaliana* (*47*). We found that the *spl2/9/10/11/13/15* mutant displayed a greatly reduced rate of DNA methylation decay in leaves that is substantially slower than in WT (Fig. 3, M to O, fig. S8B, and tables S7 and S10), consistent with a slower rate of biological aging. This reduced rate of methylation loss appears to be partially offset by young leaves emerging with a lower initial level of methylation (Fig. S8C). Together, these observations from the *miR156/SPL* pathway indicate that the rate of epigenetic aging can distinguish divergent aging rates between individuals in *A. thaliana*.

There is a wide variation in the lifespan, aging rates and DNA methylation landscapes across *A. thaliana* accessions (*53*). Thus, we tested whether epigenetic aging reflects lifespan using closely related accessions Wassilewskija-0 (Ws-0) and Ws-2, which exhibit accelerated and delayed senescence and flowering compared to WT Col-0 (grown in LD conditions), respectively (Fig. S9, A and B). We generated and analyzed methylomes for the first true leaves from various aging stages of Ws-2 and Ws-0 (Table S1) and observed an age-related mCG decay over TEs in both accessions (Fig. S9, C to E, and tables S7 and S10) that was consistent with their divergent lifespans and senescence phenotypes. Similar to *spl2/9/10/11/13/15*, the slow rate of epigenetic decay in Ws-0 was partially offset by lower levels of methylation at leaf emergence (Fig. S8D and table S7), which may be a feature of slow-aging genotypes. Together, these data demonstrate that the rate of mCG decay facilitates accurate inference of an underlying biological aging rate in *A. thaliana*, and that outliers with slow and fast aging phenotypes can be captured by DNA methylation profiling.

### The epigenetic age of individual organs is decoupled from organism age

In contrast to mammals, which typically complete organogenesis as embryos, plants form new organs continuously throughout their life cycles. This provided a unique opportunity to decouple organ age from organism age, by profiling newly emerged organs in older individuals. We generated methylomes for the first cauline leaves of WT at 8, 18, 30, and 45 days after bolting (DAB), corresponding to approximately 36, 46, 58, and 73 DAG (Table S1). Cauline leaves are a distinct leaf type that forms after the vegetative:reproductive transition (Fig. 4A). Similar to first true leaves, the first cauline leaves showed an age-related mCG decay over TEs (Fig. 4, B and C), but not genes (Fig. S10A). Young cauline leaves at 8-DAB (∼36-DAG) exhibited a “young” level of mCG in heterochromatin comparable to 18-DAG first true leaves (Fig. 4D and table S7), strongly suggesting that patterns of epigenetic decay represent organ age, rather than organism age in plants. We also confirmed this by testing newly emerged rosette leaves from 45-DAG WT plants (Fig. S10B and table S1), as well as published methylomes of cauline leaves (Table S1), which also showed mCG levels typical of young first true leaves (Fig. S10C and table S7). These findings suggest that new organs are epigenetically young and can age at independent rates from the rest of the organism. Differences in methylation levels between tissues (Fig. S10D and table S2) could therefore be influenced by age differences at the time of sampling.

**Fig. 4.**
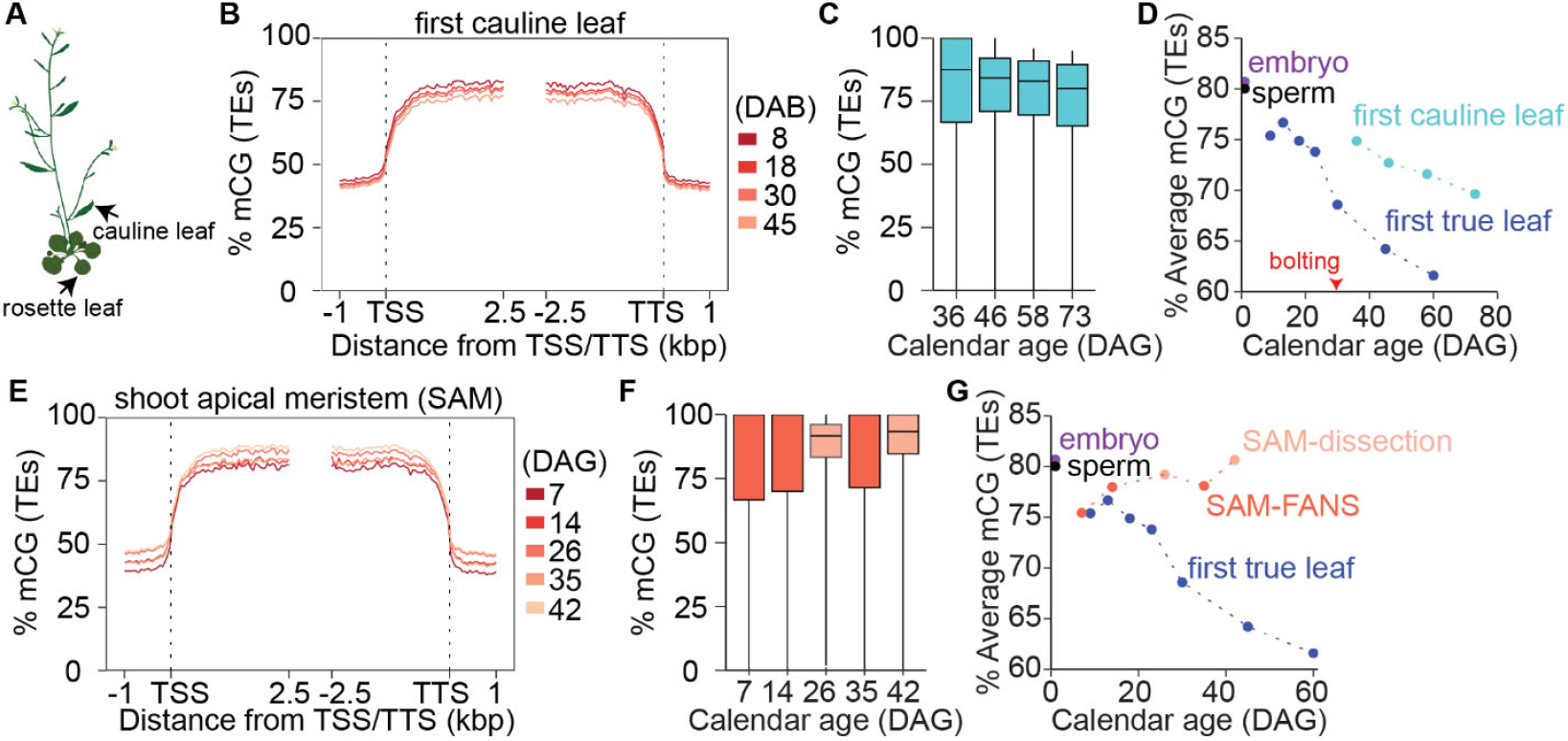
The epigenetic age of individual organs is decoupled from organism age. **(A)** Diagram annotating rosette and cauline leaves. **(B)** Distribution of mean % mCG over TEs and the flanking 1-kbp in aging first cauline leaves. **(C)** Boxplot showing the distribution of mCG in aging first cauline leaves. Boxes represent the interquartile range, horizontal lines denote the median, and whiskers denote the 10th and 90th percentiles. **(D)** Comparison of average mCG level over TEs between first true leaves and first cauline leaves. **(E)** Distribution of mean % mCG over TEs and the flanking 1-kbp in the shoot apical meristem (SAM) cells. **(F)** Boxplot showing the distribution of mCG in aging SAM. **(G)** Comparison of average mCG level over TEs between first true leaves and SAM, isolated by fluorescence-activated nuclei sorting (SAM-FANS) or manual dissection (SAM-dissection).

Plant organs initiate from stem cells within meristems. To confirm whether meristem cells are subjected to DNA methylation aging dynamics, we analyzed public methylomes of fluorescence-activated nuclear sorted (FANS) shoot apical meristem (SAM) cells from 7-, 14-, and 35-DAG plants (*54*), and dissected shoot apices from 26-(*55*) and 42-DAG (*56*) (Table S2). In contrast to the age-related mCG reduction observed in leaves, the mCG level showed no evidence of epigenetic decay, rather a slight increase in DNA methylation with age (Fig. 3, E to G, and table S7). Interestingly, the slight increase was also evident in the hematopoietic stem cell from old mice (*57*), suggesting shared features may underpin stem cell methylation dynamics during plants and animal aging. Overall, we found that SAM cells maintain a high mCG level in heterochromatin during aging comparable to embryo (Fig. 3G and table S7). These data support a model in which meristems are “ageless”, giving rise to the germline each generation, as well as somatic organs which each embark upon an independent aging program toward senescence.

### A gene regulatory program drives epigenetic aging

Currently, the molecular and developmental mechanisms underpinning DNA methylation clocks in mammals are largely unknown (*13*). To identify possible regulatory mechanisms underpinning epigenetic aging in *A. thaliana*, we profiled transcriptomes of WT first true leaves over an aging-time course (Table S4). We found that a large portion of the genome exhibits age-dependent differences in expression, culminating in 12,536 differentially expressed genes (DEGs) in old (40-DAG) vs. young (13-DAG) first true leaves (Fig. 5A, fig. S11A, and table S11). The contribution of the aging methylome to age-related transcriptome dynamics seems negligible, as the majority of DMRs are enriched in gene-poor heterochromatin (Fig. 5B and fig. S11, B and C), and only 88 DEGs (0.007% of the total) are located within 2kb of an old vs. young leaf DMR (Fig. S11D). We observed that genes underpinning the major components of DNA methylation maintenance pathways were identified as differentially expressed between young and old leaves, exhibiting a smooth decline in expression from the youngest leaves to later ages (Fig. 5C and table S12). A recent study identified that two transcriptional factors, TCX5 & TCX6, which share sequence homology with the human DREAM complex protein LIN54 (Fig. S12A), transcriptionally repress DNA methylation maintenance genes in plants (*55*). We observed that both *TCX5* and *TCX6* exhibit elevated expression during leaf aging, along with a number of additional DREAM complex homologs (Fig. S12B and table S13). We therefore hypothesized that repression of the DNA methylation maintenance machinery by TCX5/6 or a DREAM-like complex could underpin DNA methylation losses occurring during organ aging. To test this, we profiled the methylomes of the first true leaves of *tcx5/6* mutants (Fig. 5D, fig S12, C and D, and table S1), as well as several other double mutants of DREAM complex homologs (Fig. S13, A to C, and table S1). Strikingly, *tcx5/6* mutants exhibited a complete absence of epigenetic aging, maintaining high levels of DNA methylation akin to the germline, embryos, or shoot apical meristem cells (Fig. 5, E to G, fig. S12E, and table S7). This is the first example of a genetic mutant in which the DNA methylation dynamics associated with aging have been completely abolished.

**Fig. 5.**
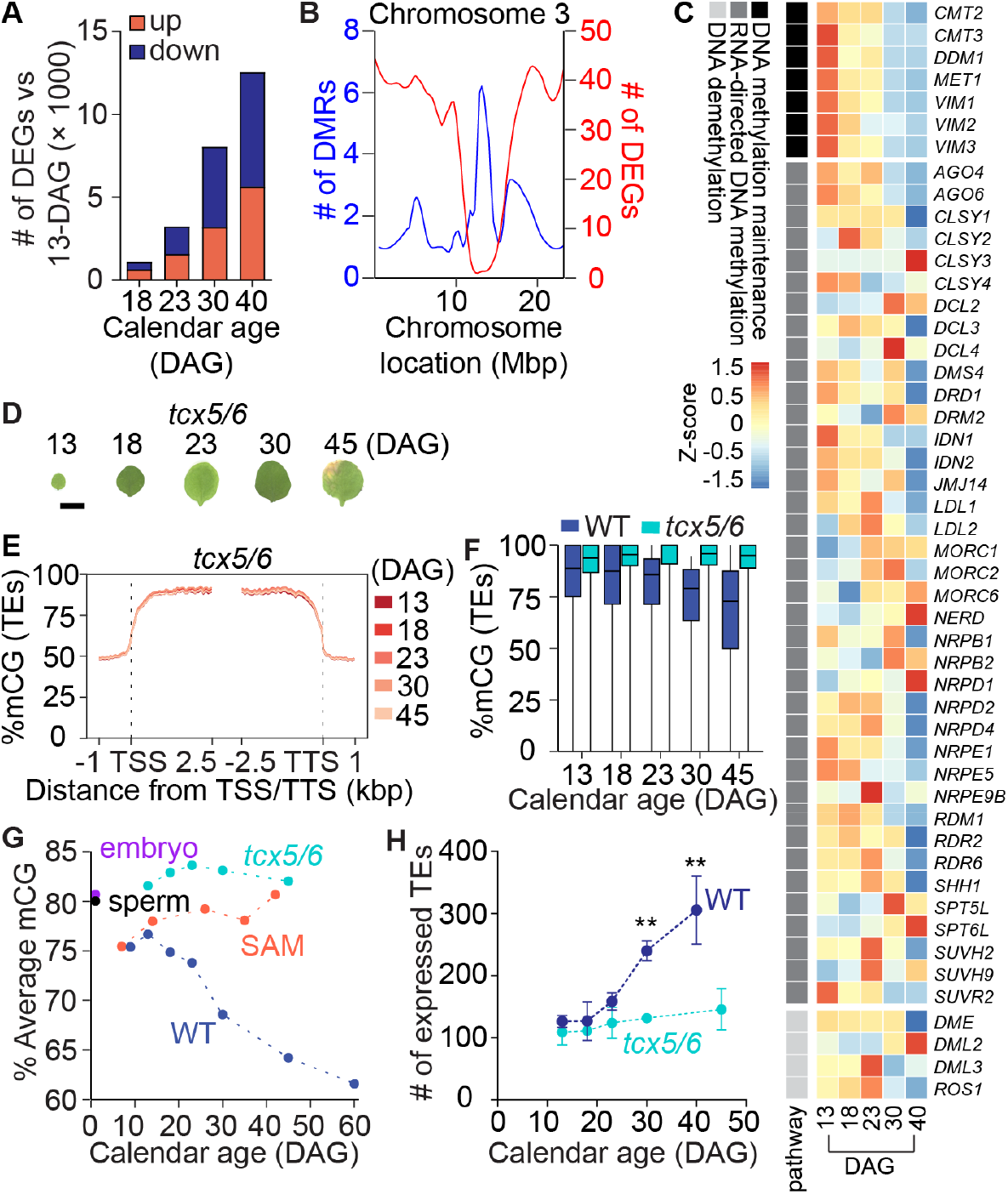
Transcriptional repression of DNA methylation maintenance genes underpins epigenetic aging. Number of DEGs in 18-, 23-, 30-, and 40-DAG WT first true leaves compared to 13-DAG. Distribution of all DEGs and DMRs compared to 13-DAG first true leaves across 250-kbp windows on chromosome 3. **(C)** Heatmap showing expression patterns of DNA methylation pathway genes. Heatmap values are Z-scores. **(D)** First true leaf phenotypes of *tcx5/6*. Bar = 0.5 cm. **(E)** Distribution of mean % mCG over TEs and the flanking 1-kbp in *tcx5/6* aging first true leaves. **(F)** Boxplot showing comparisons of mCG level in aging first true leaves between WT and *tcx5/6*. Boxes denote interquartile range, horizontal lines denote the median, and whiskers denote the 10th and 90th percentiles. All pairwise differences between WT and *tcx5/6* or mutants are significant (Mann-Whitney U test, p = <0.0001) **(G)** Comparison of average mCG level over TEs in aging first true leaves between WT and *tcx5/6*. **(H)** The number of expressed TEs during aging in first true leaves of WT and *tcx5/6*. 40-DAG WT was compared with 45-DAG *tcx5/6* for statistical analysis. Statistical significance was determined using one-tailed Welch’s t-test. **, p < 0.01.

Transcriptome profiling of *tcx5/6* mutant leaves of different ages showed that the majority of DNA methylation maintenance genes (e.g. *MET1, DDM1, VIM1*) exhibited substantially higher expression levels than WT (Fig. S12F and table S14). TCX5 binds the promoters of all these genes *in vivo* (*55*), and loss of transcriptional repression likely explains the persistence of epigenetic integrity in older *tcx5/6* organs. We also observed a moderately slower rate of epigenetic aging in *lin37a/b* mutants, but not *aly1/2* or *aly1/3* mutants (Fig. S13D). These data suggest that TCX5/6 may function as transcriptional repressors in a distinct manner to the canonical DREAM complex in animals. *tcx5/6* mutants also exhibit elevated CHG methylation (Fig. S12E), together with elevated *CMT3* expression (Fig. S12F and table S14), consistent with previous reports (*55*), whereas CHH methylation and the expression of CHH methyltransferases is unchanged (Fig. S12, E and F, and table S14). The total knockout of epigenetic aging decay in *tcx5/6* mutants is the first example of a transcriptional regulatory program required for epigenetic aging clocks.

To test if the loss of epigenetic aging may have consequences for genome regulation, we analyzed the expression of TEs in *tcx5/6* mutant vs. WT aging first true leaves. Unlike WT, *tcx5/6* mutant leaves showed no age-dependent elevation in the number of expressed TEs (Fig. 5H and table S15), suggesting that the program of epigenetic decay driven by TCX5/6 has consequences for genome defense and stability. As the expression of TEs is associated with age-influenced diseases such as cancer, the discovery of a gene regulatory program that undermines TE silencing in older tissues could have broad implications for how genome defense is maintained in aging tissues. Lastly, while *tcx5/6* mutants display moderately delayed phenotypic aging (*55*) (Fig. 5D), the overall phenotypic trajectory of aging in these mutants is fairly similar to WT, yet completely decoupled from the rate of DNA methylation loss. This finding is therefore inconsistent with provocative claims that epigenetic changes could be one of the major causes, rather than consequences, of biological aging processes in mammals (*58, 59*).

## Discussion and outlook

Our study demonstrates that a key feature of epigenetic dynamics during aging in mammals – the loss of DNA methylation within heterochromatin – is shared in the model plant *A. thaliana*. The discovery of a short-lived model species for epigenetics research with a process of age-induced epigenetic decay offers promising opportunities to gain mechanistic insights into how epigenetic aging clocks function, and whether age-induced epigenetic changes cause genomic instability in a manner that could impact disease. Additionally, the discovery that plant shoot apical meristems do not exhibit epigenetic aging may lead to new discoveries into how a regularly dividing population of cells can maintain high epigenetic fidelity over long periods of time. This also likely explains why long-lived perennial tree species do not show signs of epigenetic decay in newly emerged organs (*60, 61*). Our study demonstrates that the age-related loss of epigenetic information in heterochromatin is driven by a gene regulatory program which silences DNA methylation maintenance pathways during organ aging. Understanding how this program responds to the environment, growth rate, development, senescence and physiological drivers of aging will help uncover the broader functional significance of epigenetic aging clocks in eukaryotes and whether they can be stalled or reversed.

## Supporting information

Combined_supplementary_materials

## Acknowledgments

We thank Peilei Deng (Kookmin University) for helping prepare illustration of *A. thaliana*.

## Author contributions

Conceptualization: DD & BPW. Methodology: DD, KC & BPW. Investigation: DD, KC & JT. Visualization: DD. Funding acquisition: BPW. Project administration: BPW. Supervision: DD & BPW. Writing – original draft: DD & BPW. Writing – review & editing: DD, KC & BPW

## Data and materials availability

All high throughput sequencing data will be deposited on the NCBI GEO after acceptance for publication.

## Supplementary Materials

Materials and Methods

Figs. S1 to S13

Tables S1 to S18

References (*1*–*13*)

