## Supplementary material for "Aging drives a program of DNA methylation decay in plant organs": Combined_supplementary_materials

**The PDF file includes:**

Materials and Methods
Figs. S1 to S13
References

**Other Supplementary Materials for this manuscript include the following:**

Table S1 to S18

### Materials and Methods

#### Plant materials

All mutant lines used in this study are in the Columbia-0 (Col-0) background. Ws-0 (CS22623), Ws-2 (CS22659), *aly1* (SALK_114476), *aly2* (SALK_118765C), *aly3* (SALKseq_40756), *lin37a* (SALK_057175C), *lin37b* (SALK_103139C), *tcx5* (SALK_144605C), *tcx6* (CS443483), *mir156a/b/c/d* (CS71704), *mir156a/c mir157a/c* (CS71710), *spl2/9/10/11/13/15* (CS69799), *V9A:rSPL15* (CS2109787) seeds were obtained from the Arabidopsis Biological Resource Center (ABRC) and were confirmed with subsequent genotyping. The *aly1/2*, *aly1/3*, *lin37a/b*, and *tcx5/6* mutants were generated by genetic crossing (*aly1* × *aly2*, *aly1* × *aly3*, *lin37a* × *lin37b*, *tcx5* × *tcx6*) and subsequent genotyping in F2 populations. Primer sequences are listed in Table S18.

#### Plant growth and sample collection

Seeds were stratified for 3 days (10 days for Ws-0 and Ws-2) at 4 °C in the dark and directly sown on water-saturated soil. Plants were grown either under long-day conditions (LD, 16 h light/ 8 h dark, 22 °C, 60% humidity) in a Percival growth chamber (AR100L3), or under short-day conditions (SD, 8 h light/ 16 h dark, room temperature) in a temperature-controlled greenhouse.

For EM-seq purpose, LD-exposed first true leaves of Col-0 at 9-, 13-, 18-, 23-, 30-, 45-, and 60-days after gemination (DAG), Ws-0, Ws-2, *mir156a/b/c/d*, *mir156a/c mir157a/c*, *spl2/9/10/11/13/15*, *V9A:rSPL15*, *and tcx5/6* at 13-, 18-, 23-, 30-, and 45-DAG, *aly1/2*, *aly1/3*, and *lin37a/b* at 13-, 18-, 23-, and 30-DAG were collected. LD-exposed first cauline leaves of Col-0 at 8-, 18-, 30-, and 45-days after bolting (DAB) were collected. Around 50 mature green embryos of Ws-2 and *spl2/9/10/11/13/15* were dissected from plants grown under LD conditions. For plants grown under SD, the first true leaves of Col-0 at 13-, 18-, 23-, 30-, and 45-DAG were collected.

For quantitative RT-PCR and RNA-seq purposes, the first true leaves of Col-0 at 13-, 18-, 23-, 30-, and 40-DAG, *tcx5/6* at 13-, 18-, 23-, 30-, and 45-DAG were collected from plants grown under LD conditions with three biological replicates. All leaf samples were collected at about 2-3 pm of each date to avoid circadian effects. All collected tissues were immediately frozen in liquid nitrogen and kept at −80 °C until use.

#### Enzymatic methyl sequencing

DNA was isolated using the CTAB method. Briefly, plant tissues were ground using a Qiagen TissueLyser II and treated with CTAB buffer (300 µl) at 65 °C for approximately 30 min. RNA was removed by incubating with 3 µl of RNase (Thermo Fisher Scientific) at 37 °C for 20 min. This was followed by the addition of 300 µl chloroform, 10-min centrifugation and ethanol washes to isolate total genomic DNA. The DNA concentration was quantified using a Qubit Flex fluorometer. DNA samples were sonicated to an average fragment size of ~300 bp using a Covaris S220 Focused Ultrasonicator (99-s treatment, 140 peak power, 10 duty factor, and 200 cycles/burst). EM-seq libraries were constructed following the NEBNext® Enzymatic Methyl-seq Kit protocol with 50 ng initial DNA (25 ng for all embryo samples). Library quality was verified with a KAPA Library Quant Kit and fragment analyzer. Libraries were pooled and sequenced using an Illumina NextSeq 2000 (150 bp, paired-end reads) or an NovaSeq X (150 bp, paired-end reads). Further information on each EM-seq library sequenced in this study is available in Table S1.

**Whole-genome methylation analysis**

A total of 133 published methylomes were obtained and analyzed. Mapping of whole-genome bisulfite sequencing and EM-seq data was performed as described (*1*). Briefly, reads were preprocessed with TrimGalore 0.6.6 (Babraham Bioinformatics) to remove adaptors. Bismark 0.22.3 (*2*) was used to map filtered reads to the *A. thaliana* TAIR10 genome, allowing for 0 mismatch per read in the seed region and removing PCR duplicates. Methylation values for each cytosine were calculated as #C/(#C+#T) using the Bismark methylation extractor function. The efficiency of EM-seq conversion was verified by quantifying the percentage of methylation for reads mapped to the chloroplast. EM-seq conversion rates were > 99.4% across all samples sequenced in this study. For analysis of DNA methylation patterns in various tissues and shoot apical meristem cells, biological replicates were pooled prior to mapping. A summary of all published methylomes used in this study is presented in Table S2.

mCG difference was calculated by comparing methylation levels of individual cytosines in aging leaf samples against embryo. Negative and positive values were called hypo- and hyper-cytosines, respectively. DMRs were identified using the R package DMRcaller 1.36.0 (*3*). Briefly, the genome was divided into 300 bp bins containing at least 10 cytosines with >= 5 reads coverage per cytosine. Filtered bins with an absolute CG methylation difference of at least 0.35 and cutoffs of Benjamini–Hochberg corrected false discovery rate < 0.01 were computed as DMRs.

**Quantitative RT-PCR and RNA-seq**

Total RNA was isolated from the first true leaves of Col-0 and *tcx5/6* at 13-, 18, 23-, 30-, and 40-DAG (45-DAG for *tcx5/6*) grown under LD using a QIAGEN RNeasy plant mini kit. For qRT-PCR analysis, 100 ng total RNA were reverse-transcribed with HiScript III RT Supermix (Vazyme) and cDNA was amplified at target loci using Fast SYBR™ Green Master Mix (Thermo Fisher Scientific) and a QuantStudio 3 qPCR system (Applied Biosystems). The constitutively expressed *TIP41* (AT4G34270) was used as an endogenous control for normalization (*4*). All reactions were performed in three biological replicates, with two technical replicates for each reaction. Raw Ct values and primer sequences are listed in Table S17 and Table S18, respectively.

For RNA-seq, non-stranded libraries from poly-A-tailed RNA were generated according to the manufacturer's instructions (Illumina) and sequenced on an NovaSeq 6000 using 150-bp paired-end protocol performed by Novogene. Three biological replicates were used for all RNA-seq experiments. Mapping was performed as previously described (*1*). Briefly, adapters were trimmed using cutadapt (*5*) and Trim Galore 0.6.6 (Babraham Bioinformatics) by trimming 9 bp from the 5'-end of reads and enforcing a 3'-end quality of > 25%. Reads were mapped to the Araport11 genome using STAR 2.7.1a (*6*), permitting 0.05 mismatches as a fraction of total read length and discarding reads that did not map uniquely. Normalization of reads count was performed by running htseq-count of HTSeq 2.0 (*7*) and DESeq2 1.40.1 (*8*). Normalized reads count of each library is available in Table S16. Differentially expressed genes were identified based on a minimum of two-fold change in expression and a Benjamini–Hochberg corrected P-value < 0.05. To identify expressed TEs, read counts for all annotated full length transposable elements (annotations from Araport 11) were collated from htseq-count. 6 out of 3901TEs that were constitutively and highly expressed in all samples were excluded as potential domesticated TEs. The number of expressed TEs was calculated by counting all TEs with >0 reads for each sample. Further information on each RNA-seq library sequenced in this study is available in Table S4. A summary of all published transcriptomes used in this study is presented in Table S5.

### Associating DMRs with proximal genes

All DEGs and DMRs from pairwise comparisons between aging (18-, 23-, 30-, and 40-45-DAG) and young (13-DAG) leaf samples were pooled for association analysis. DEGs located within 2-kbp of a DMR were extracted. Genome-wide distribution of all DEGs and DMRs was plotted within 250-kbp bins in R.

### Elastic net regression

R package Caret v. 6.0.94 (*9*) was used to train a predictive elastic net regression model on 101 methylomes from newly generated and published data. The 101 methylomes were split into training (80 samples) and testing (21 samples) sets (chosen at random) (Table S8). CpG sites were retained for training if they were present in at least 90% of training samples with read depth >=5, using median imputation to fill in missing or low read depth methylation values. To train an elastic net (EN) regression model, we tested all values at 0.1 increments between 0 and 1 values for the relative weight of the ridge and lasso penalties (alpha) and all values between 0.0001 and 4 at 0.1 increments for the overall strength of the regularization (lambda). Minimizing root mean squared error (RMSE) after evaluation using 5-times repeated 5-fold validation resulted in a final model (alpha = 0.1 and lambda = 1.4) that identified 276 CG sites as high confidence predictors for aging. List of 276 CG sites was available in Table S9.

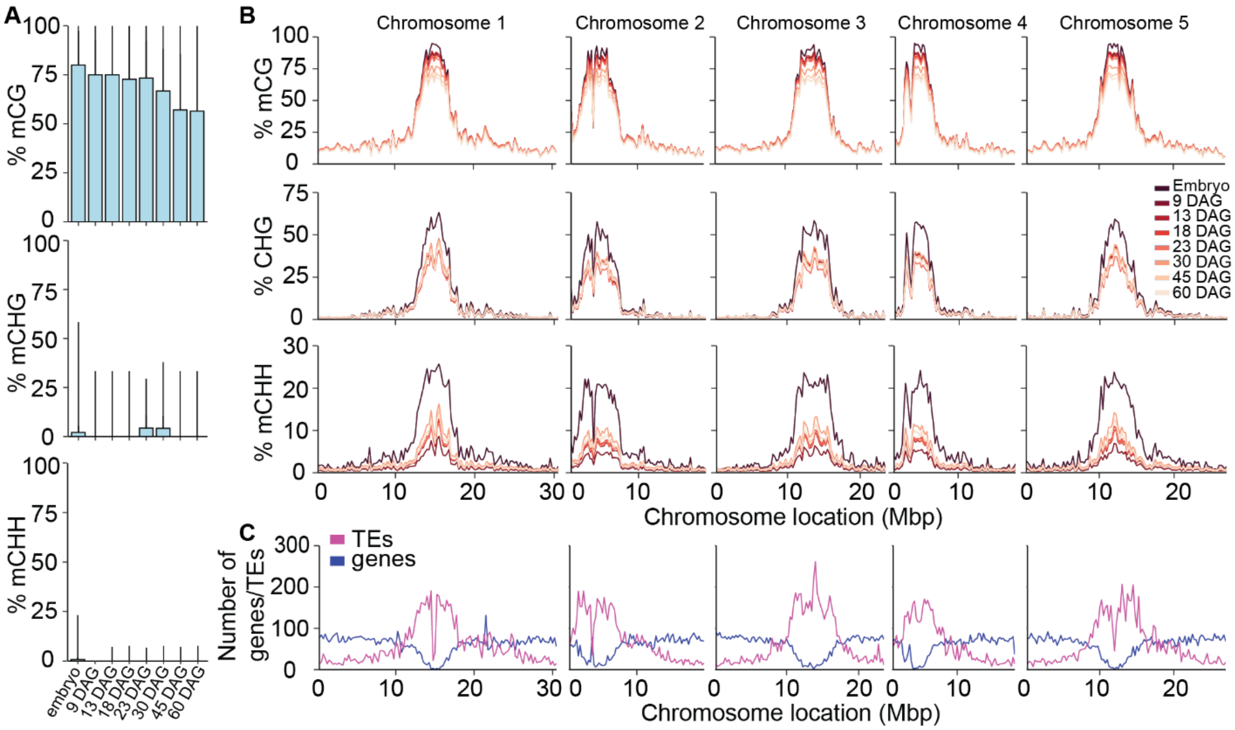


**Fig. S1. DNA methylation dynamics during leaf aging in *A. thaliana*.**

**(A)** Boxplot showing the distribution of DNA methylation in all three cytosine contexts. Boxes represent the interquartile range, horizontal lines denote the median, and whiskers denote the 10th and 90th percentiles. **(B)** Genome-wide distribution of DNA methylation in all three cytosine contexts with 250-kbp windows in embryo and aging first true leaves (WT). The methylation pattern of embryo was used as an early life reference (*10*). **(C)** Chromosomal distribution of TEs and genes.

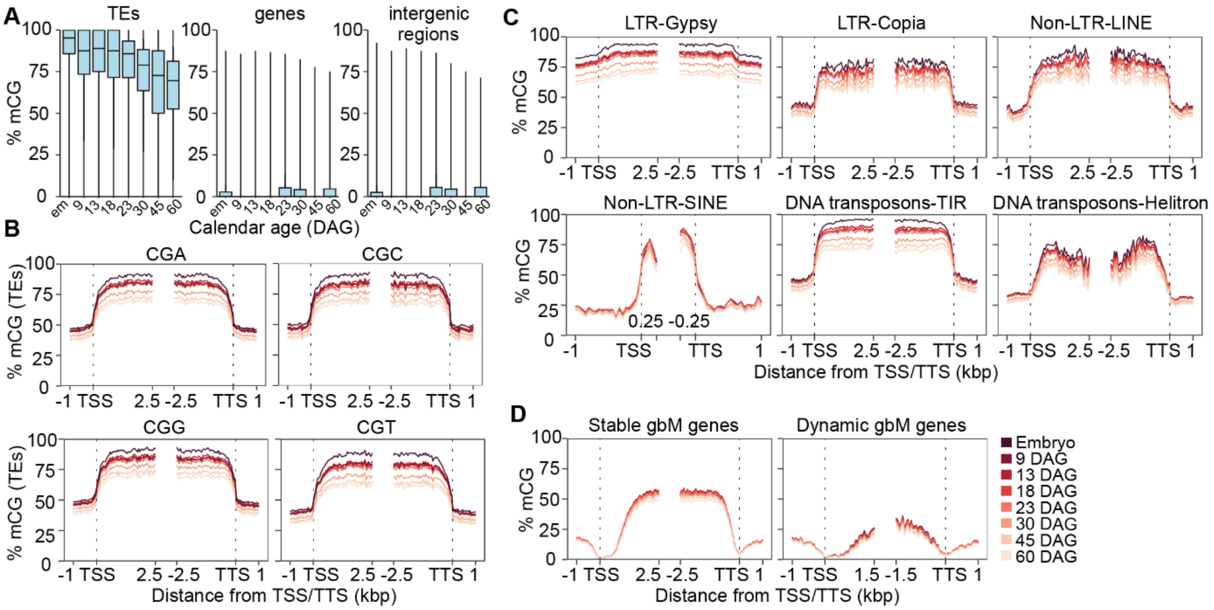


**Fig. S2. Heterochromatin mCG decay during leaf aging in *A. thaliana*.**

**(A)** Boxplot showing the distribution of mCG over TEs, genes, and intergenic regions. Boxes represent the interquartile range, horizontal lines denote the median, and whiskers denote the 10th and 90th percentiles. (**B** to **D**) Distribution of mean % mCG within TEs across all CG sub-contexts **(B)**, TE superfamilies **(C)**, and subtypes of gene body methylated (gbM) genes **(D)**.

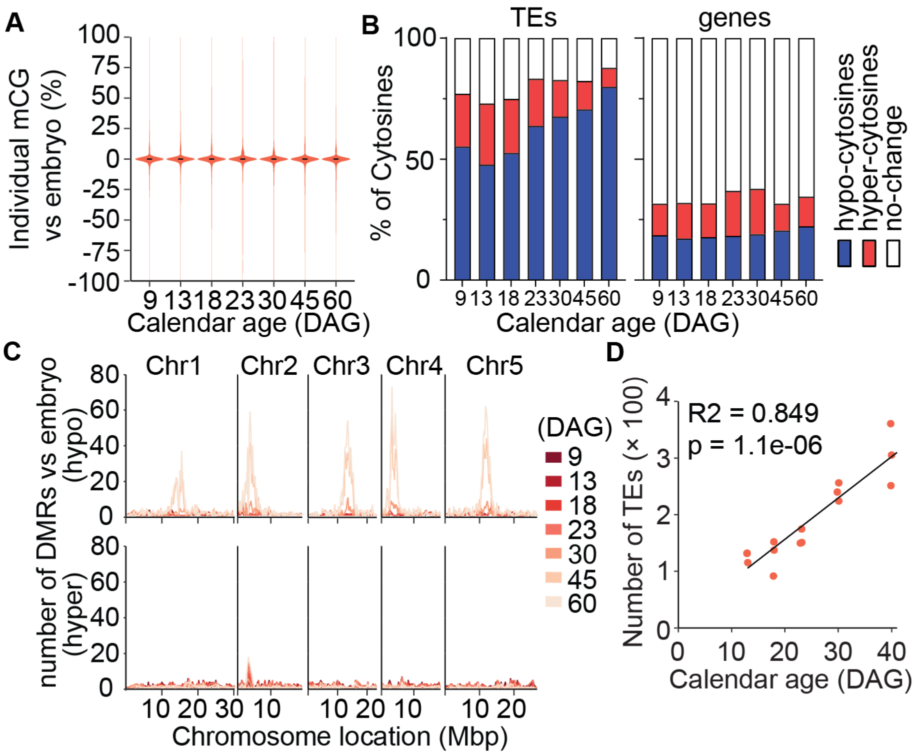


**Fig. S3. Impact of aging on individual cytosines.**

**(A)** Violin plot showing the difference in percent methylation of individual CGs (within genes) between embryo and aging first true leaves. Boxes denote the interquartile range, horizontal lines represent the median, and whiskers denote the 10th and 90th percentiles. **(B)** Proportion of cytosines (CGs only) showing reduced (hypo) and increased (hyper) methylation in aging first true leaves compared to embryo. **(C)** Chromosomal distribution of CG DMRs with 250-kbp windows in aging first true leaves compared to embryo. **(D)** The number of expressed TEs in WT first true leaves over an aging time course from our RNA-seq data.

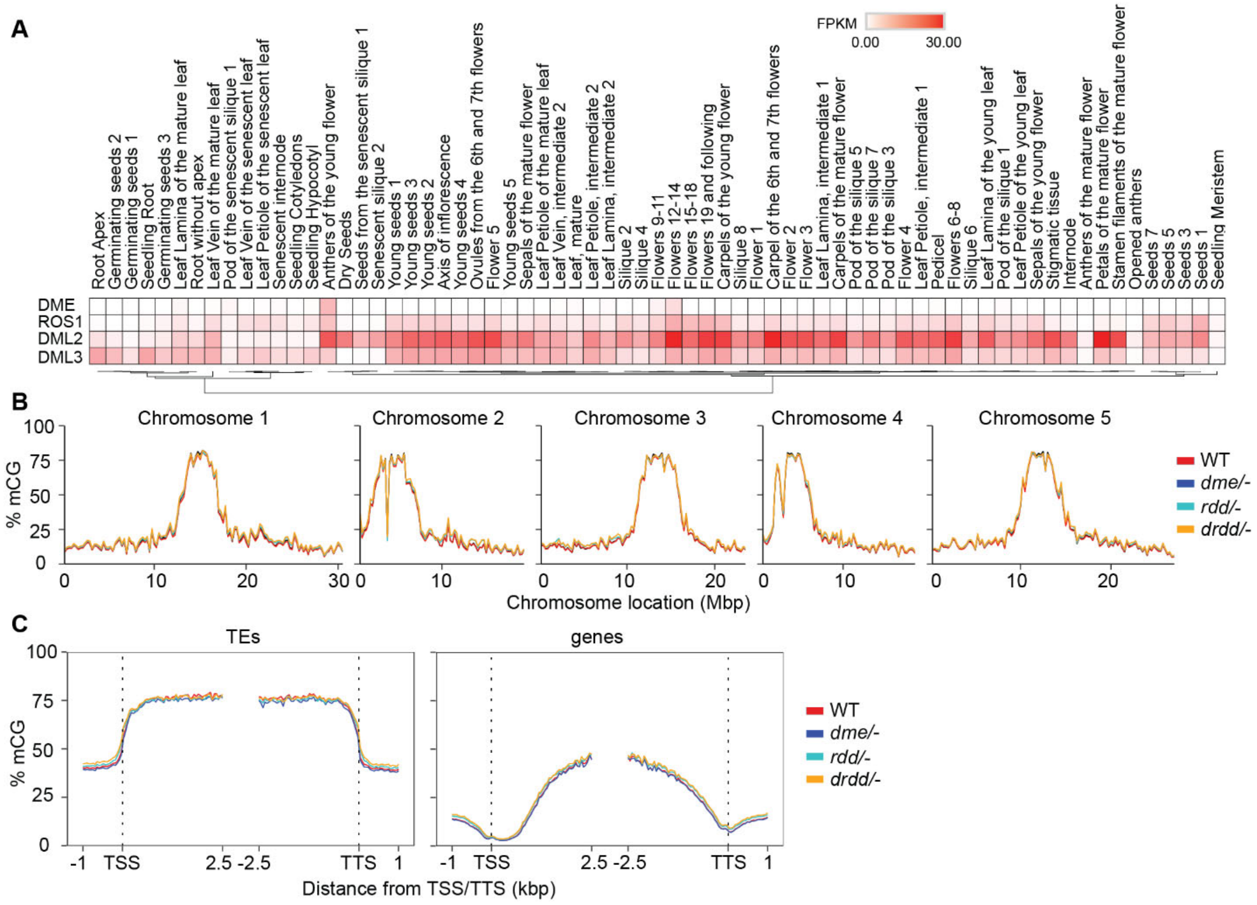


**Fig. S4. DNA demethylases do not impact age-related DNA methylation dynamics.**

**(A)** Published transcriptome data for DRDD (DME, ROS1, DML2, DML3) demethylases (*11*). The color scale indicates the relative expression level based on normalized FPKM values. **(B)** Genome-wide distribution of mCG with 250-kbp windows in 21-DAG WT (WT segregants for *drdd*), *dme*, *rdd*, and *drdd* mutant leaves (*12*). **(C)** Distribution of mean % mCG across total TEs and genes and the flanking 1-kbp regions in WT, *dme*, *rdd*, and *drdd* mutants.

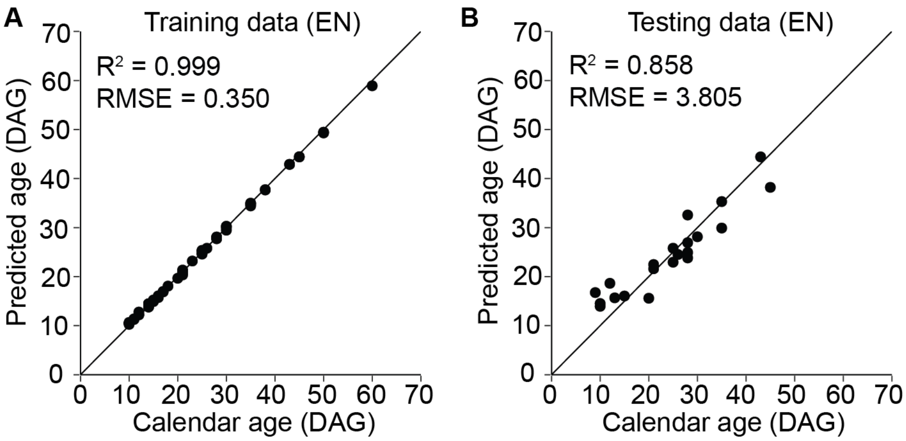


**Fig. S5. Training and testing results based on 234 EN-identified CG sites within TEs.**

**(A)** Predictive accuracy of a trained elastic net regression model on training data. **(B)** Predictive accuracy of a trained elastic net regression model on testing data.

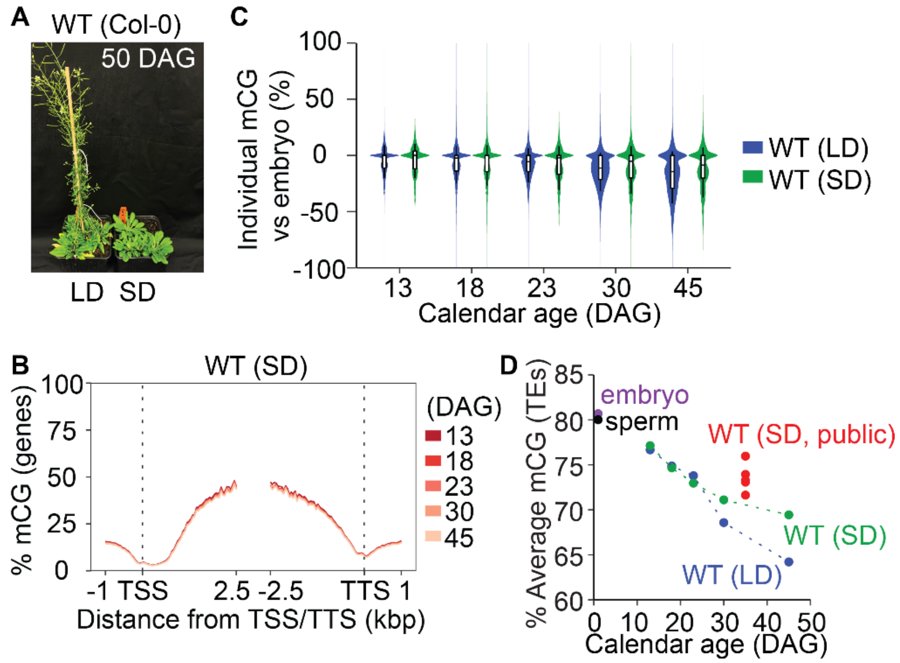


**Fig. S6. SD conditions slow down epigenetic aging.**

**(A)** 50-DAG *A. thaliana* WT plants grown in LD and SD conditions. **(B)** Distribution of mean % mCG across total genes and the flanking 1-kbp regions in SD-grown aging first true leaves. **(C)** Violin plot showing the difference in percent methylation of individual CGs (within TEs) between LD and SD conditions. Boxes bound the interquartile range, horizontal lines represent the median, and whiskers denote the 10th and 90th percentiles. **(D)** Average mCG level over TEs in aging first true leaves from LD and SD conditions. Published methylomes from SD-grown rosette leaves (not first true leaves) were obtained and analyzed. Average mCG levels over TEs in embryo (*10*) and sperm (*13*) were used as early life references.

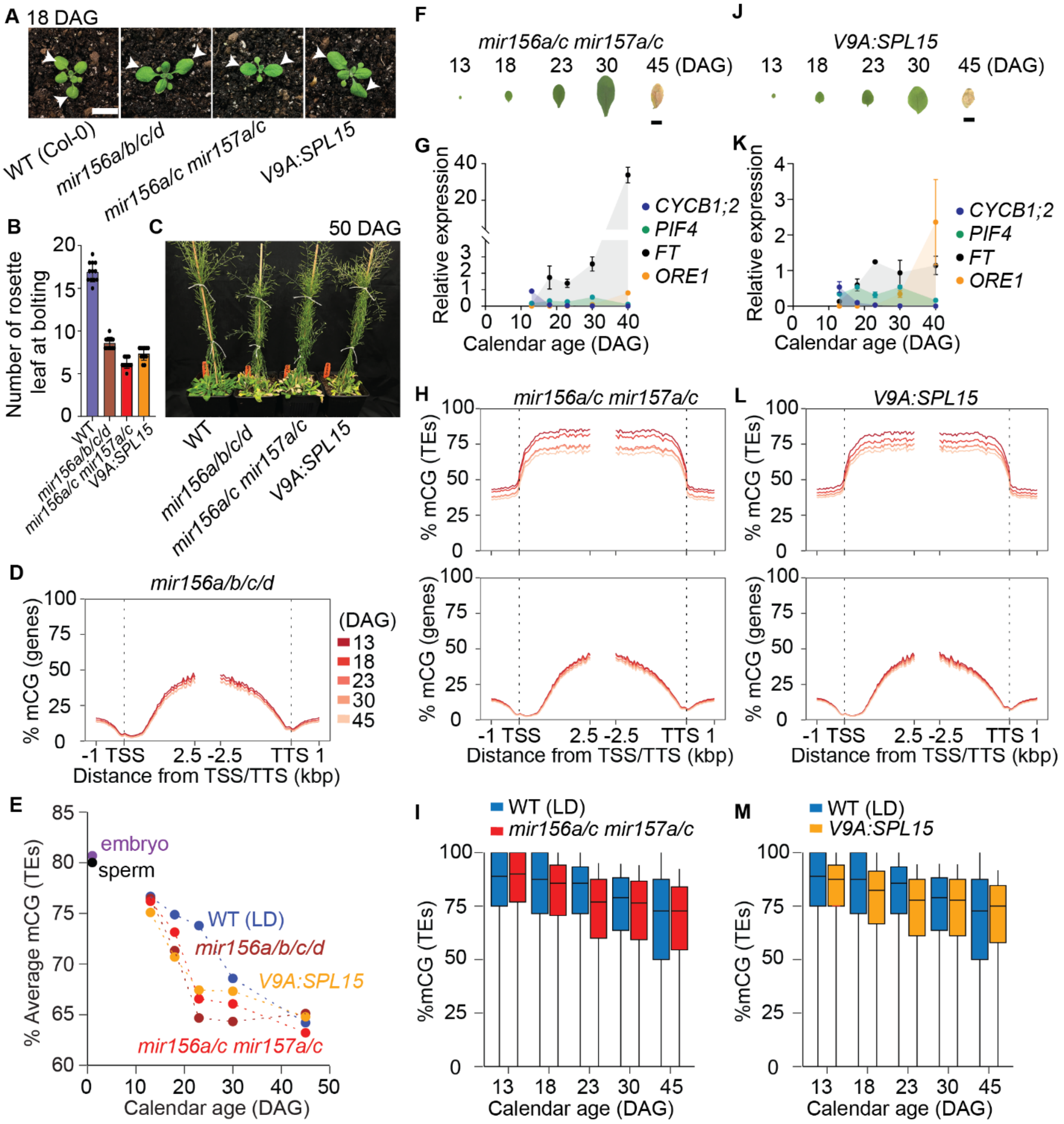


**Fig. S7. Epigenetic aging in *mir156/157* mutants.**

**(A)** 18-DAG seedlings of WT, *mir156a/b/c/d*, *mir156a/c mir157a/c*, and *V9A:rSPL15* grown in LD conditions. Arrows denote altered first true leaf morphology consistent with an accelerated juvenile:adult transition in mutants. Bar = 1 cm. **(B)** The number of rosette leaves in WT, *mir156a/b/c/d*, *mir156a/c mir157a/c*, and *V9A:rSPL15* at bolting. Values are means ± SD (n = 12). **(C)** 50-DAG plants of WT, *mir156a/b/c/d*, *mir156a/c mir157a/c*, and *V9A:rSPL15* grown in LD conditions. **(D)** Distribution of mean % mCG across genes in aging first true leaves of *mir156a/b/c/d*. **(E)** Average mCG level over TEs in aging first true leaves of WT, *mir156a/b/c/d*, *mir156a/c mir157a/c*, and *V9A:rSPL15*. (**F** and **J**) First true leaf aging phenotypes of *mir156a/c mir157a/c* (**F**) and *V9A:rSPL15* (**J**). Bars = 0.5 cm. (**G** and **K**) Quantitative RT-PCR analysis of

*CYCB1;2*, *PIF4*, *FT*, and *ORE1* transcript levels in first true leaves of *mir156a/c mir157a/c* **(G)** and *V9A:rSPL15* **(K)** from various aging stages. Error bars represent standard deviations of the mean from three biological replicates. All values of each gene were normalized relative to its highest expression in WT (LD) (set as 1). (**H** and **L**) Distribution of mean % mCG across TEs and genes and the flanking 1-kbp in *mir156a/c mir157a/c* **(H)** and *V9A:rSPL15* **(L)** aging first true leaves. (**I** and **M**) Boxplot showing the distribution of mCG over TEs in *mir156a/c mir157a/c* **(I)** and *V9A:rSPL15* **(M)** aging first true leaves. Boxes denote the interquartile range, horizontal lines denote the median and whiskers denote the 10th and 90th percentiles.

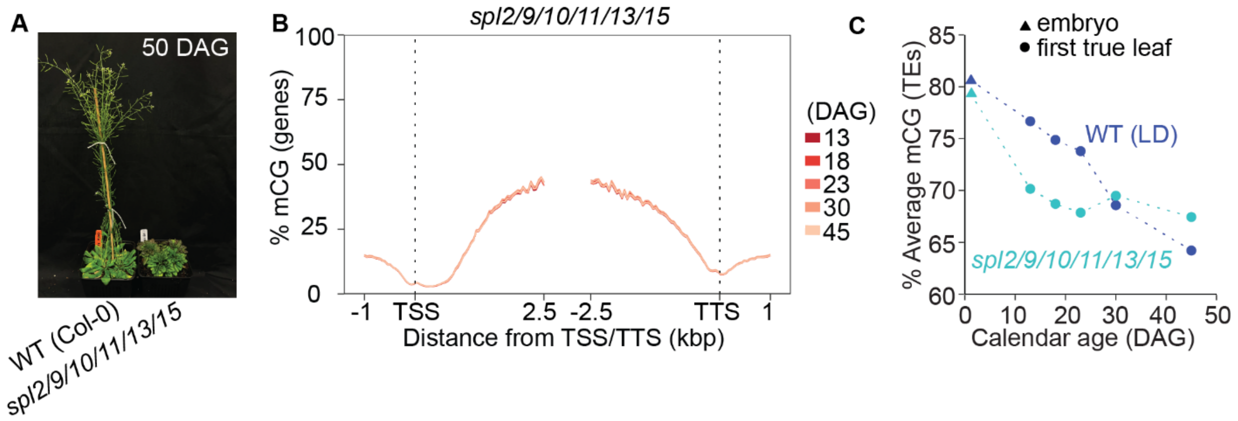


**Fig. S8. Epigenetic aging in *spl2/9/10/11/13/15* mutants. (A)** 50-DAG plants of WT and *spl2/9/10/11/13/15* grown in LD conditions. **(B)** Distribution of mean % mCG across genes in aging first true leaves of *spl2/9/10/11/13/15*. **(C)** Average mCG level over TEs in embryo and aging first true leaves of WT and *spl2/9/10/11/13/15*.

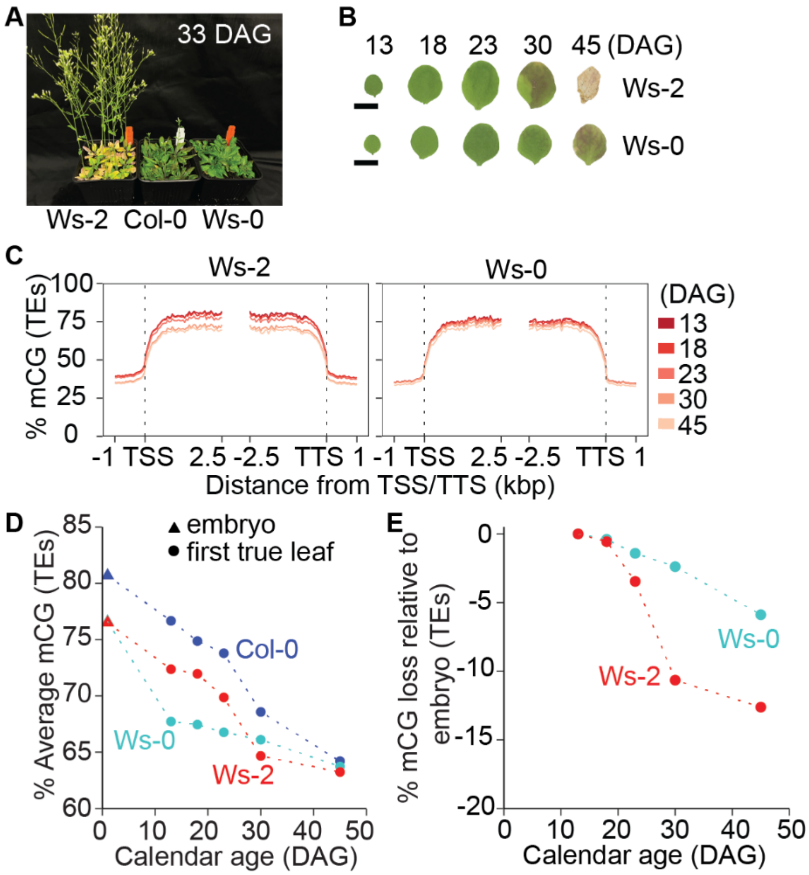


**Fig. S9. Epigenetic aging in Ws-2 and Ws-0 accessions. (A)** 33-DAG plants of WT (Col-0), Ws-2, and Ws-0 accessions grown in LD conditions. **(B)** First true leaf aging phenotypes of Ws-2 and Ws-0. Bars = 0.5 cm. **(C)** Distribution of mean % mCG across TEs in aging first true leaves of Ws-2 and Ws-0. **(D)** Average mCG level over TEs in embryo and aging first true leaves of WT (Col-0), Ws-2 and Ws-0. **(E)** The rate of mCG decay in aging first true leaves of Ws-2 and Ws-0.

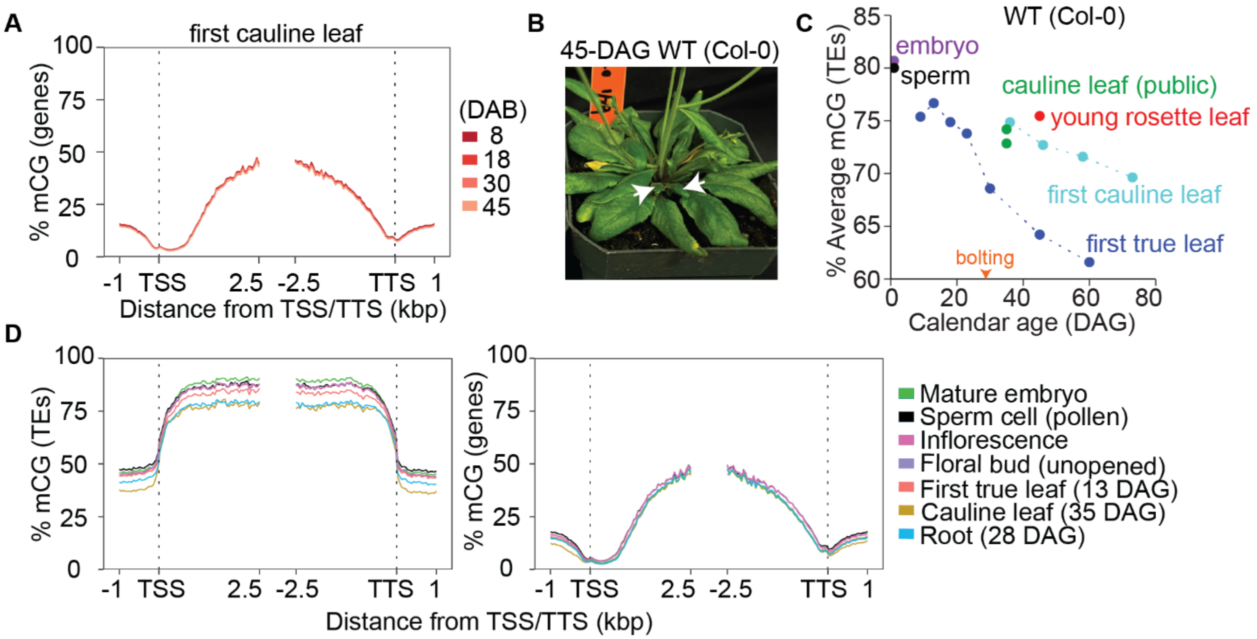


**Fig. S10. Epigenetic aging in first cauline leaves and young rosette leaves. (A)** Distribution of mean % mCG across genes in aging first cauline leaves. **(B)** 45-DAG rosette of WT. The arrows indicate rosette leaves assessed for methylation profiling. **(C)** Average mCG level over TEs in aging first true leaves, aging first cauline leaves, and 45-DAG young rosette leaves. Public methylomes of cauline leaves were obtained and analyzed. The arrow indicates the bolting stage. **(D)** Distribution of mean % mCG across TEs and genes in various WT tissues.

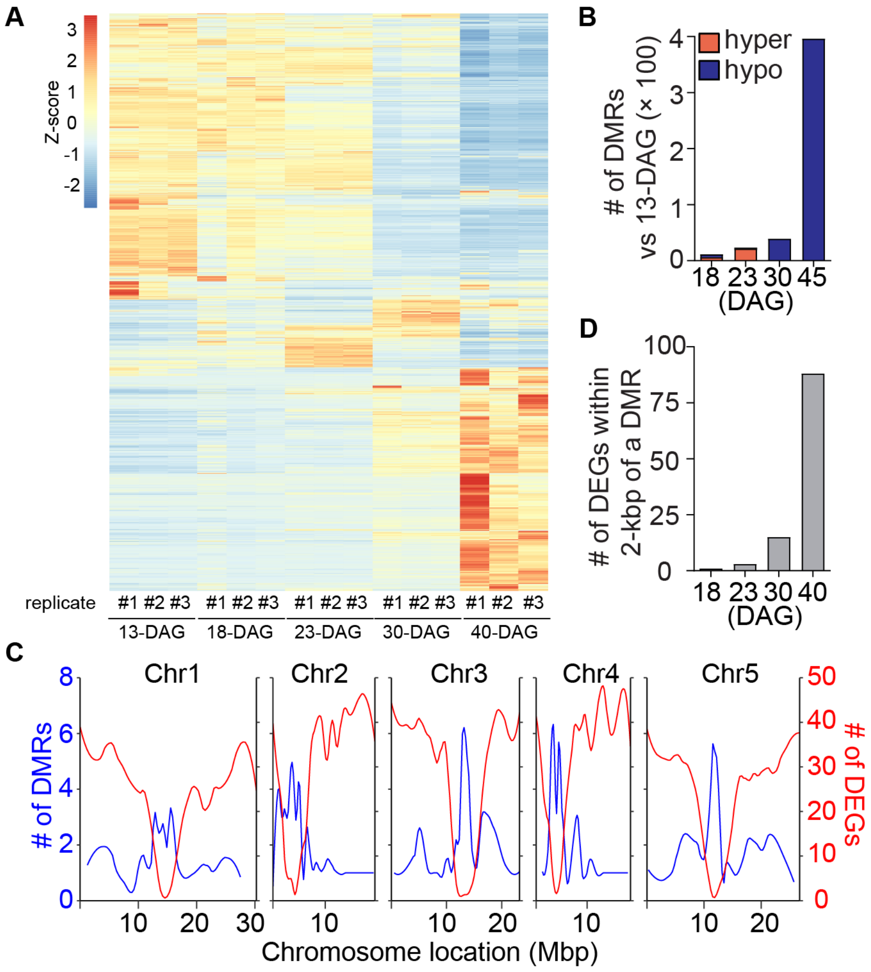


**Fig. S11. Aging transcriptome of WT first true leaves.**

**(A)** Heatmap showing expression patterns of 14175 DEGs from pairwise comparisons between 13-DAG and 18-, 23- 30-, and 40-DAG WT first true leaves. Genes with a Bonferroni adjusted p value < 0.05 and fold change >2 were defined as DEGs in pairwise comparisons. Heatmap values are Z-scores. **(B)** The number of CG DMRs in 18-, 23-. 30, and 45-DAG first true leaves compared to 13-DAG. **(C)** Chromosomal distribution with 250-kbp windows of all CG DMRs and DEGs from pairwise comparisons against 13-DAG first true leaves. **(D)** The number of DEGs within 2-kbp of a CG DMR. DMRs in 45-DAG first true leaves were used to associate DEGs in 40-DAG first true leaves.

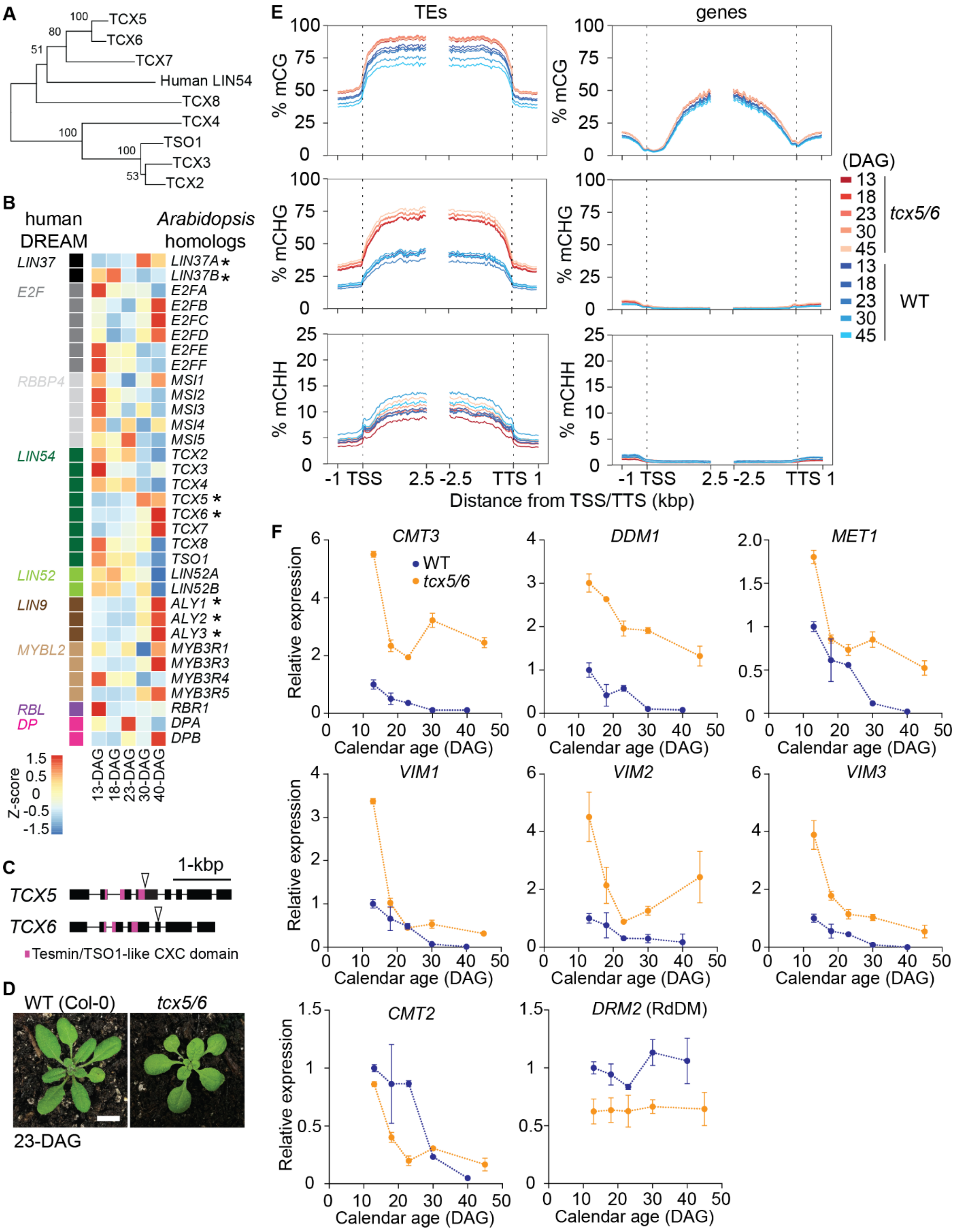


**Fig. S12. Epigenetic aging in *tcx5/6* mutants.**

**(A)** Gene tree showing evolutionary relationship between human *LIN54* and *Arabidopsis TCX* genes. **(B)** Heatmap showing expression patterns of *A. thaliana* homologs of human DREAM complex components in aging WT first true leaves. Heatmap values are Z-scores. **(C)** Gene diagrams of T-DNA mutant alleles used in this study. T-DNA insertions are marked by triangles. Black boxes are exons, solid lines are introns. (**D**) 23-DAG seedling phenotypes of WT and *tcx5/6* grown under LD conditions. Bar = 1 cm. **(E)** Distribution of mean % mCG across TEs and genes in WT and *tcx5/6* aging first true leaves. **(F)** Expression patterns of genes involved in DNA methylation maintenance in WT and *tcx5/6* aging first true leaves. Error bars represent standard deviations of the mean from three biological replicates. All values of each gene were normalized relative to its expression at 13-DAG in WT (set as 1).

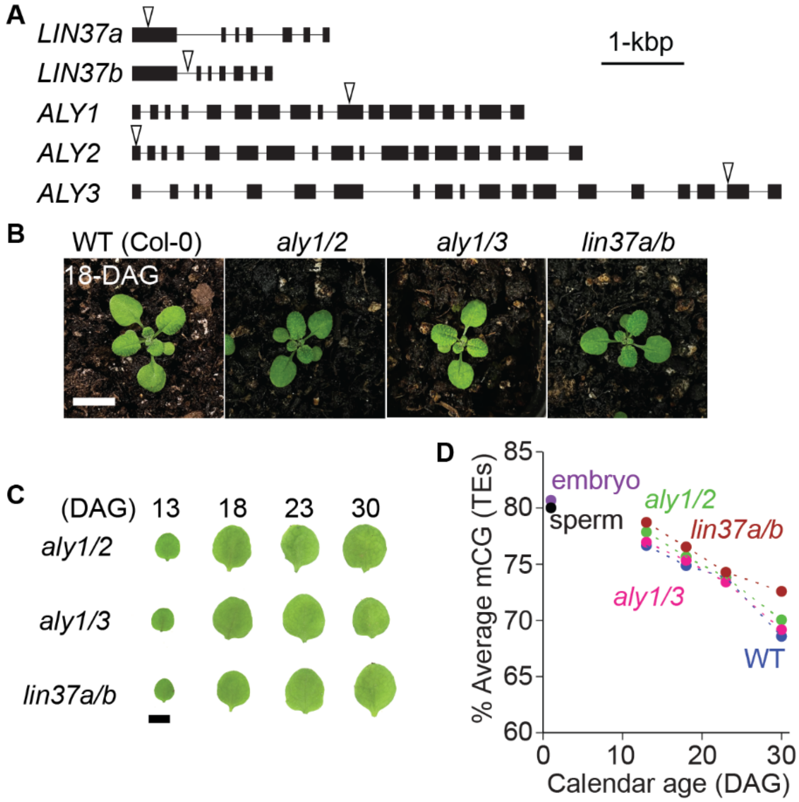


**Fig. S13. Epigenetic aging in *aly1/2*, *aly1/3*, and *lin37a/b* mutants.**

**(A)** Gene diagrams of T-DNA mutant alleles used in this study. T-DNA insertions are marked by triangles. Black boxes are exons, solid lines are introns. (**D**) 18-DAG seedling phenotypes of WT, *aly1/2*, *aly1/3*, and *lin37a/b* grown under LD conditions. Bar = 1cm. **(C)** First true leaf aging phenotypes of *aly1/2*, *aly1/3*, and *lin37a/b*. Bar = 0.5 cm. **(D)** Average mCG level over TEs in aging first true leaves of WT, *aly1/2*, *aly1/3*, and *lin37a/b*.
